# Biofilms deform soft surfaces and disrupt epithelia

**DOI:** 10.1101/2020.01.29.923060

**Authors:** Alice Cont, Tamara Rossy, Zainebe Al-Mayyah, Alexandre Persat

## Abstract

During chronic infections and in microbiota, bacteria predominantly colonize their hosts as multicellular structures called biofilms. Despite their ubiquity *in vivo*, we still lack a basic understanding of how they interact with biological tissues, and ultimately how they influence host physiology. A common assumption is that biofilms interact with their hosts biochemically. However, the contributions of mechanics, while being central to the process of biofilm formation, have been vastly overlooked as a factor influencing host physiology. Specifically, how biofilms form on soft, tissue-like materials remains unknown. Here we show that biofilms can deform soft substrates by transmission of internally-generated mechanical stresses. We found that biofilms from both *Vibrio cholerae* and *Pseudomonas aeruginosa* can induce large deformations of soft synthetic hydrogels. Using a combination of mechanical measurements and mutants in matrix components, we found that biofilms deform their substrates by simultaneous buckling and adhesion. Specifically, mechanical constraints opposing growth causes biofilm buckling, while matrix components maintaining surface adhesion transmit buckling forces to the substrate. Finally, we demonstrate that biofilms can generate sufficient mechanical stress to deform and disrupt soft epithelial cell monolayers, suggesting that these forces can damage a host independently of typical virulence factors. Altogether, our results illustrate that forces generated by bacterial communities play an important role not only in biofilm morphogenesis but also in host physiology, suggesting a mechanical mode of infection.

## Introduction

In their natural environments, bacteria commonly grow and self-organize into multicellular structures called biofilms (1). Biofilms form when bacteria attach onto a solid surface and divide while embedding themselves in a matrix of extracellular polymeric substances (EPS) (2). The biofilm matrix is a viscoelastic material generally composed of a mixture of polysaccharides, proteins, nucleic acids and cellular debris (3). EPS maintains cell-cell cohesion throughout the lifecycle of a biofilm, also making the resident cells more resilient to selective pressures. The biofilm lifestyle provides resident cells with fitness advantages compared to their planktonic counterpart, for example by increasing their tolerance to external chemical stressors such as antimicrobials and host immune effectors. In addition, its mechanical strength and cohesion promotes biofilm integrity against physical challenge such as flow and grazing (4).

Bacteria commonly colonize the tissues of their host in the form of biofilms. For example, biofilms are a common contributor of infections, for example in cystic fibrosis patients who are chronically infected by biofilms of the opportunistic pathogen *P. aeruginosa* (5, 6). Biofilms are also widespread in microbiota, for example as commensals seek to stably associate to host intestinal epithelium (7). As they grow on or within a host, biofilms must cope with a battery of chemical and physical stressors. In particular, they must inevitably form at the surface of soft biological material composed of host cells or extracellular matrix (ECM). Despite host-associated biofilms ubiquitously forming on soft surface, we still lack a rigorous understanding of how the mechanical properties of a substrate impacts the physiology of a biofilm, and reciprocally how biofilms impact the mechanics of soft biological surfaces.

The growth of single cells embedded within self-secreted EPS drives biofilm formation. During this process, cells locally stretch or compress the elastic matrix, thereby exerting mechanical stress (8, 9). This local action at the level of single cells collectively generates mechanical stress across the whole biofilm structure. Thus, the combination of biofilm growth and matrix elastic properties imposes the generation of internal mechanical stress (10). As a consequence of this stress, bacterial colony biofilms form folds and wrinkles when growing on agar plates or at an air-liquid interface (11, 12). These mechanics also influences the spatial organization of single cells within *V. cholerae* immersed biofilms (13, 14). Internal mechanical stress can also arise by a combination of cell-surface adhesion and growth, influencing the architecture of submerged biofilms and microcolonies. Friction force between the microcolony and the surface opposes biofilm expansion, generating an inward internal stress that leads to a buckling instability verticalizing or reorienting contiguous cells (14, 15). These studies demonstrate the importance of mechanics in biofilm morphogenesis and spatial organization, but their function in the context of host colonization remains unknown.

Here, we investigate how biofilms form at the surface of soft material whose mechanical properties replicate the ones encountered *in vivo*. We show that biofilms from the model pathogens *V. cholerae* and *P. aeruginosa* can deform soft synthetic hydrogel substrates they grow on. By spatially and quantitatively measuring substrate morphology, we propose a model where biofilms buckle to initiate deformations. Using EPS matrix mutants we demonstrate that deformations of the substrate require EPS matrix components maintaining cell-cell cohesion and cell-surface adhesion. The magnitude of the deformations depends on the stiffness of the material in a range that is consistent with host properties. Using traction force microscopy, we show that biofilms can generate large mechanical stress in the MPa range. Finally, we demonstrate that biofilms can deform and even damage tissue-engineered soft epithelia whose mechanics reproduce the ones of a host-tissue. These insights suggest that forces generated by growing biofilms could play a role not only in biofilm morphomechanics, but also in mechanically compromising the physiology of their host.

## Results

### Biofilms deform soft substrates

To understand how biofilms interact with soft surfaces, we first explored their formation on synthetic hydrogel substrates. We generated polyethylene glycol (PEG) hydrogel films via photoinitiated polymerization of PEG diacrylate precursors at the bottom surface of microfluidic channels. These polymeric films are covalently bound to the glass surface to avoid drift and delamination. By using a “sandwich” method for polymerization, we could fabricate flat ~100 µm-thin PEG films that allowed us to perform high resolution live confocal imaging of biofilm formation under flow (Fig. 1A). We used the *V. cholerae* A1152 strain (*V. cholerae* WT*) which constitutively produces large amounts of EPS matrix, thereby generating robust and reproducible biofilms. On soft hydrogels, *V. cholerae* formed biofilms whose bottom surfaces appeared bell-shaped (Fig. 1*B*), in striking difference with the typically flat-bottom biofilms that form on hard surfaces such as glass and plastic. To distinguish whether this shape was a result of the deformation of the hydrogel or of the detachment of the biofilm from the surface, we embedded fluorescent tracer particles within the hydrogel film by mixing them with the pre-polymer solution before the cross-linking step. We could observe that the fluorescent tracer particles filled the apparent bell-shaped void at the biofilm core and that the hydrogel surface and the biofilm remained in contact (Fig. 1*C*). This demonstrates that the soft hydrogel substrate deforms under *V. cholerae* biofilms.

**Fig. 1:**
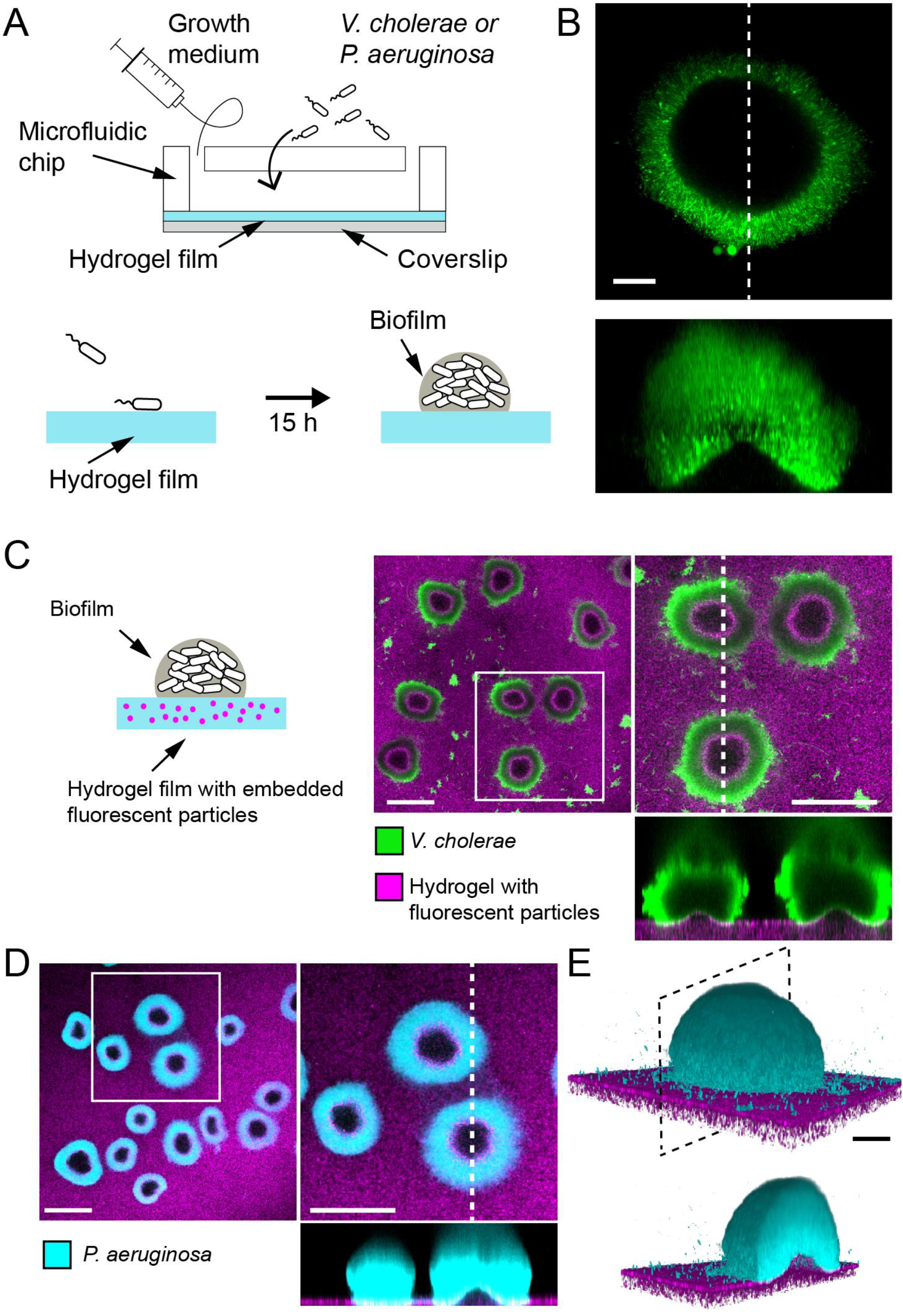
Biofilms deform soft substrates. (A) Illustration of experimental setup where we generate thin hydrogel films at the bottom surface of microchannels. These devices allow us to study biofilm formation on hydrogels reproducing mechanical properties of host tissues. (B) In-plane and cross-sectional confocal visualizations show that *V. cholerae* biofilms growing on hydrogels display large gaps at their core. (C) Embedding fluorescence tracer particle in the hydrogel films allow for visualization of deformations. *V. cholerae* biofilms formed at the surface of the films deform the substrate. (D) *P. aeruginosa* biofilms similarly deform the soft substrates. Hydrogel elastic modulus: (B and C) *E* = 12 kPa, (D and E) *E* = 38 kPa. Scale bars: (C and D) 100 µm, (B and E) 20 µm.

We then wondered whether these deformations were specifically induced by *V. cholerae* or could represent a common feature of biofilms across species. To answer this, we tested whether *P. aeruginosa* biofilms could deform soft hydrogels. We found that biofilms of *P. aeruginosa wspF^−^* mutant (*P. aeruginosa* WT*), which constitutively produces large amounts of EPS matrix, could similarly deform soft PEG hydrogels (Fig. 1*D-E*), and so did wild-type (Fig. S1). In summary, *V. cholerae* and *P. aeruginosa*, two model biofilm-forming species with distinct EPS composition are both able to deform soft substrates. This is consistent with a mechanism where biofilms generate mechanical stress on the material they grow on.

### Biofilm deform soft substrates after reaching a critical diameter

How could biofilms mechanically deform their substrates? Given the influence of growth-induced internal mechanical stress on biofilm morphology and architecture, we hypothesized that biofilms could deform soft substrates by transmission of internal stresses to the substrate they grow on. To test this hypothesis, we performed dynamic visualizations of the deformations of the hydrogel film as biofilms grew. To obtain an accurate deformation profile, we performed a radial re-slicing and averaging around the biofilm center. We could thus extract the deformation profile *δ*, its maximum deformation amplitude *δ*_*max*_ and full-width at half maximum *λ* (Fig. 2*A*). We thus recorded surface profiles for many biofilms. By reconstructing hydrogel surfaces for biofilms of different sizes, we found that *δ*_*max*_ and *λ* linearly scaled with the diameter *d* of the biofilm (Fig. S2), indicating that biofilm expansion promotes surface deformation.

**Fig. 2:**
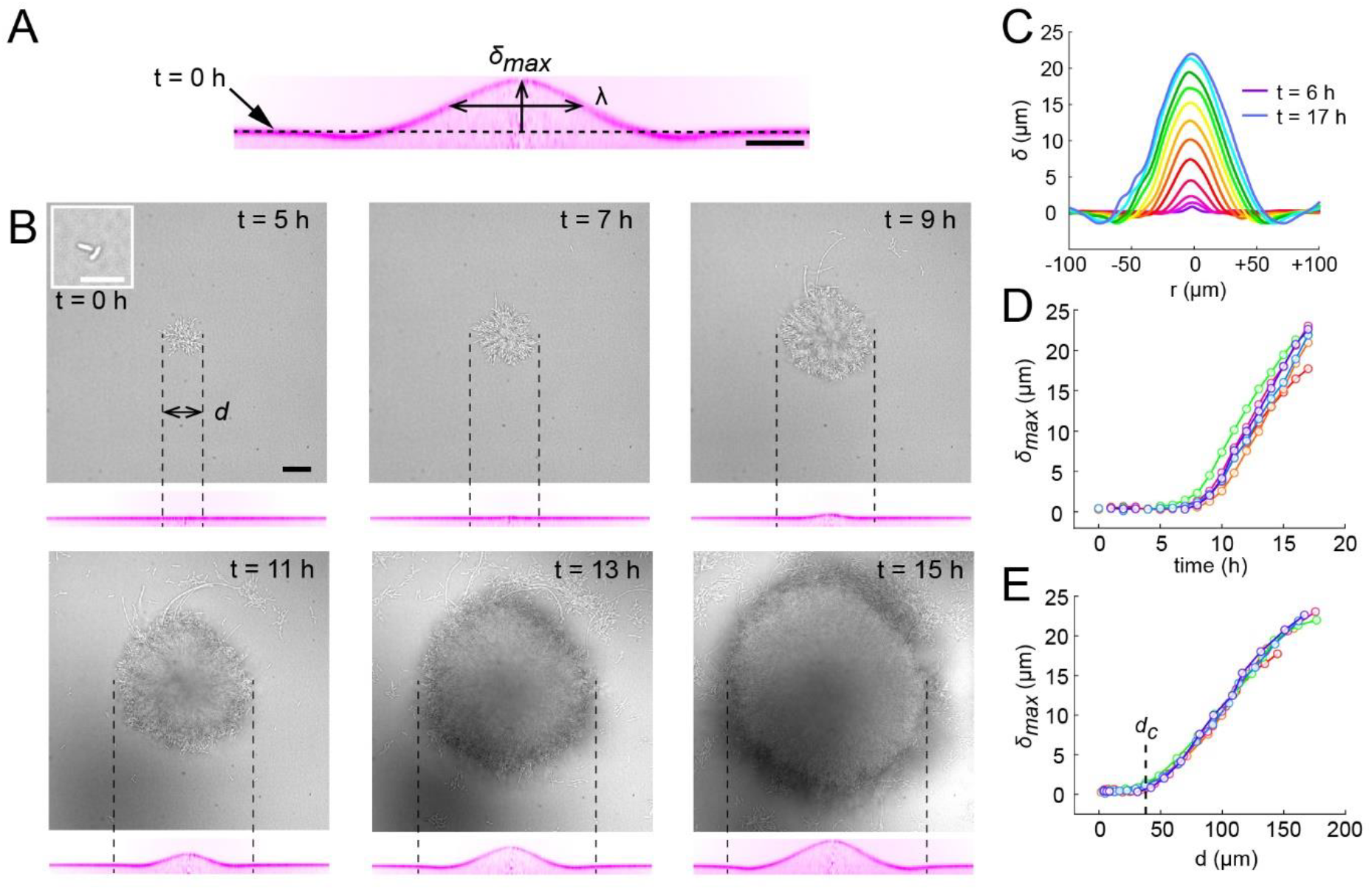
Substrate deformation dynamics highlight a critical biofilm diameter. (A) Morphological parameters *δ*_*max*_ (maximum deformation amplitude) and *λ* (half max full width) computed from resliced deformation profiles. Dashed line indicates the baseline position of the gel surface. (B) Timelapse visualization of *V. cholerae* biofilm growth (brightfield, top) with deformation (reslice, bottom). Dashed lines indicate biofilm position and size on the corresponding hydrogel profile. (C) Superimposition of these profiles shows the rapid deformation and the emergence of a recess at biofilm edges. Each color corresponds to the same biofilm at different times. (D) Time evolution of *δ*_*max*_ shows a rapid increase after 6 to 7 h of growth. (E) The dependence of *δ*_*max*_ on biofilm diameter highlights a critical biofilm diameter *d*_*c*_ above which deformation occurs. For D and E each line color corresponds to a different biofilm. Scale bar: 10 µm for inset *t* = 0 h in (B), else 20 µm.

We went further and dynamically tracked these deformations for single biofilms. Deformations increased as biofilms grew, even displaying a slight recess near the biofilm edges (Fig. 2*B-C*, Movie S1). In these visualizations, we noticed that there was a lag between the increase in biofilm diameter and the onset of deformation, with a finite deformation only appearing after 7 h of growth. This was further confirmed by following the deformations generated by many biofilms. Measurable morphological changes of the surface appeared after 6 to 7 h of growth (Fig. 2*D*). Rescaling these measurements with the diameter of the biofilm collapsed *δ*_*max*_ measurements, highlighting a critical biofilm diameter (35 µm) above which deformations emerged (Fig. 2*E*). The existence of a critical diameter is reminiscent to buckling instabilities of rigid bodies subject to compressive stress, as in Euler buckling.

### Biofilms push their substrate in the growth direction

To further investigate the mechanism by which biofilms deform surfaces, we quantified the hydrogel substrate strain during growth. To achieve this, we tracked the displacements of the fluorescent tracer particles embedded within the hydrogel in 3D using a digital volume correlation algorithm (16). At the early stages of hydrogel deformation, we found that in the plane defined by the initial surface at rest, the particles under the biofilm move in the direction of growth. Thus, the strain field shows that the biofilm stretches its substrate radially in the outward direction in addition to vertical deformations (Fig. 3*A* and Fig. S3). In other words, a biofilm applies an in-plane stress on the substrate in its growth direction, which is most likely generated by a friction between the biofilm and the surface (14, 15). As a result, the elastic biofilm experiences a force in the opposite direction, towards its center. In summary, the opposition between biofilm growth and friction with the surface generates an internal mechanical stress within the biofilm oriented radially, towards its center.

**Fig. 3:**
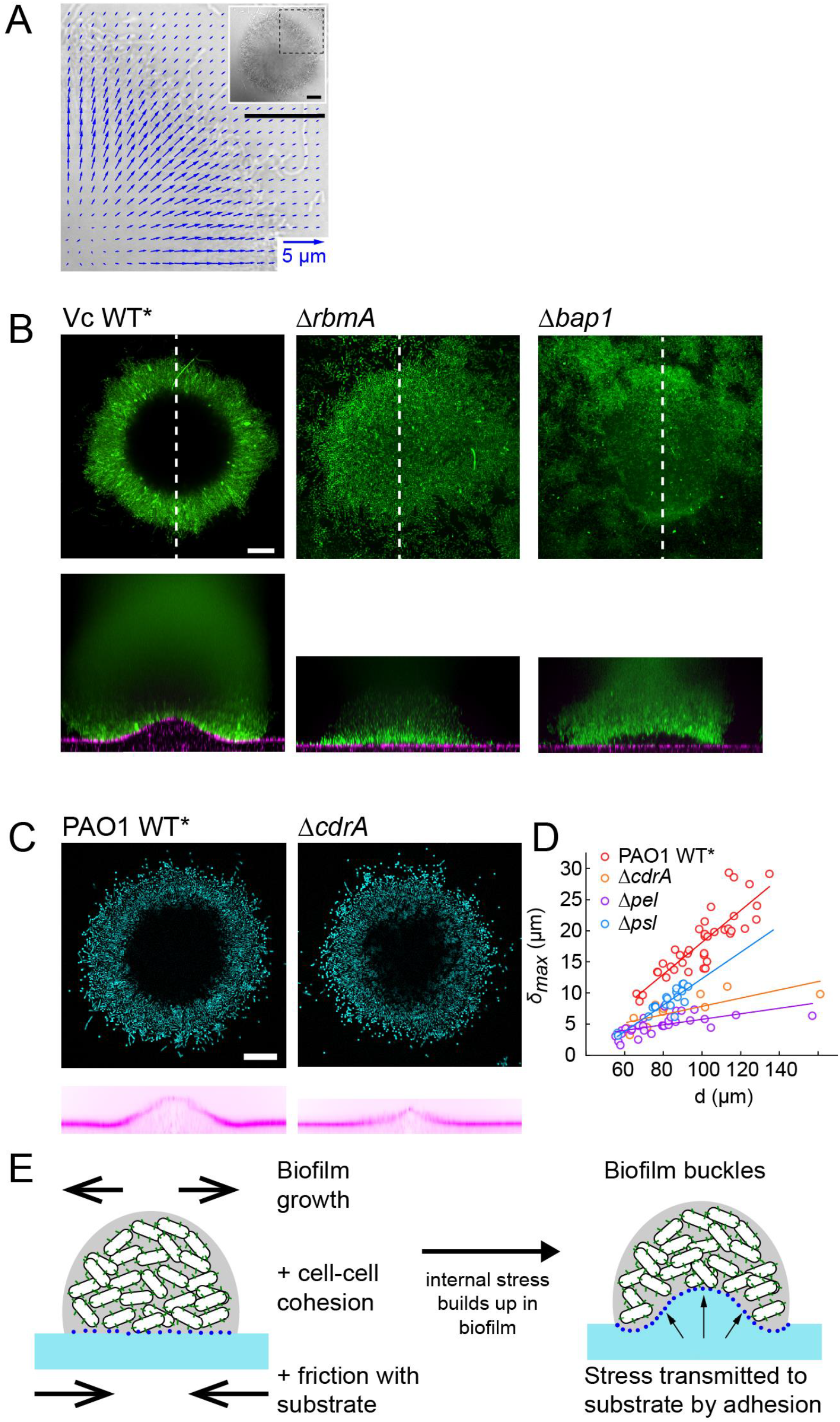
Biofilms deform their growth substrate by buckling. (A) Hydrogel strain field computed by digital volume correlation between 11 h and 12 h of growth. We superimposed the vector strain field with a brightfield image of the biofilm. For visualization purposes e only display data for the top right quarter of the biofilm shown in inset (dashed lines). (B) Deformations of hydrogel substrates by *V. cholerae* WT*, *rbma*^−^ and *bap1*^−^ biofilms. Biofilms formed by *rbma*^−^ and *bap1*^−^ fail to deform the substrate. *bap1*^−^ biofilms delaminate from the hydrogel surface. (C) Comparison of hydrogel deformations by *P. aeruginosa* WT* and *cdrA*^−^ biofilms. (D) Dependence of maximum deformations on *P. aeruginosa* WT*, *cdrA*^−^, *pel*^−^ and *psl*^−^ biofilm diameter. All matrix mutants tend to generate weaker deformations compared to WT*. (E) A model for the mechanism of biofilm deformation of soft substrates. Buildup of mechanical stress in the biofilm induces buckling. Adhesion between the biofilm and the surface transmits buckling-generated stress to the hydrogel, inducing deformations. Scale bars: 20 µm.

### EPS drives biofilm and substrate deformations

We then wondered how mechanical properties of biofilms influence substrate deformations. To investigate their contributions, we used *V. cholerae* EPS matrix mutants with altered biofilm structure and mechanical properties. The *V. cholerae* matrix is mainly composed of a polysaccharide (*vps*) and proteins including Rbma, an extracellular component which specifically strengthens cell-cell cohesion and stiffens the matrix (17, 18). We found that biofilms of *rbmA* deletion mutants were unable to deform the hydrogel substrate, demonstrating that cell-cell cohesion is an essential ingredient in force generation (Fig. 3*B*). In *P. aeruginosa*, the polysaccharides Pel and Psl, and the protein CdrA play partially redundant functions in maintaining elastic properties of the biofilm (19–21). In a similar manner, we found that the deformations generated by *P. aeruginosa* mutants in these matrix components are decreased compared to WT*, but are not abolished (Fig. 3*C*). Specifically, deletion mutants in *psl, pel* and *cdrA* showed a decrease in deformation amplitude, further demonstrating that mechanical cohesion plays a key role in surface deformation (Fig. 3*C-D*). We observed the strongest decrease in deformation for deletion mutants in *pel*.

We then probed the function of adhesion of the biofilm with the surface by visualizing the deformations generated by a *V.cholerae bap1* deletion mutant. Bap1 is specifically secreted at the biofilm-substrate interface to maintain proper surface attachment (18). The *bap1*^−^ mutant formed biofilms that did not deform the surface. However, it produced biofilms that were slightly bent but which delaminated from the substrate, thereby creating a gap between the biofilm and the hydrogel, indicating that it may have buckled (Fig. 3*B*). Our observations of the *bap1* mutant show that adhesion transmits mechanical stress generated by buckling from the biofilm to the substrate. Due to the redundant functions of its EPS components, we could not produce *P. aeruginosa* mutants with altered surface adhesion properties. However, *P. aeruginosa* biofilms growing on hydrogels with large Young’s modulus delaminated. This highlight that the transition between deformed and delaminated substrate depends on the relative contribution of adhesion strength and substrate elasticity (Fig. S6). In summary, cell-cell mechanical cohesion is essential in generating the internal stress that promotes biofilm buckling, while cell-substrate adhesion transmits this stress to the underlying substrate (Fig. 3*E*).

### Biofilms generate large traction forces

Biofilms thus deform soft materials by combining of growth-induced buckling and adhesion to their substrate. Could the mechanical stress generated on the substrate also impact various types of biological surfaces? To first explore this possibility, we quantified the forces exerted by the biofilm on hydrogel films. We used our previous particle tracking data to perform traction force microscopy, thereby computing the stress field and surface forces applied by the biofilm on the hydrogel. Traction forces were surprisingly large, reaching 5 MPa at the biofilm center after 12 h of growth (Fig. 4*A*). We note that the magnitude of the stress is relatively large, reaching the value of typical turgor pressure which in essence drives biofilms growth and stretching (22). In comparison, epithelial cell-cell junctions break when experiencing a few kPa (23). Therefore, we anticipate that biofilms produce sufficient force to mechanically deform and potentially dismantle epithelia.

**Fig. 4:**
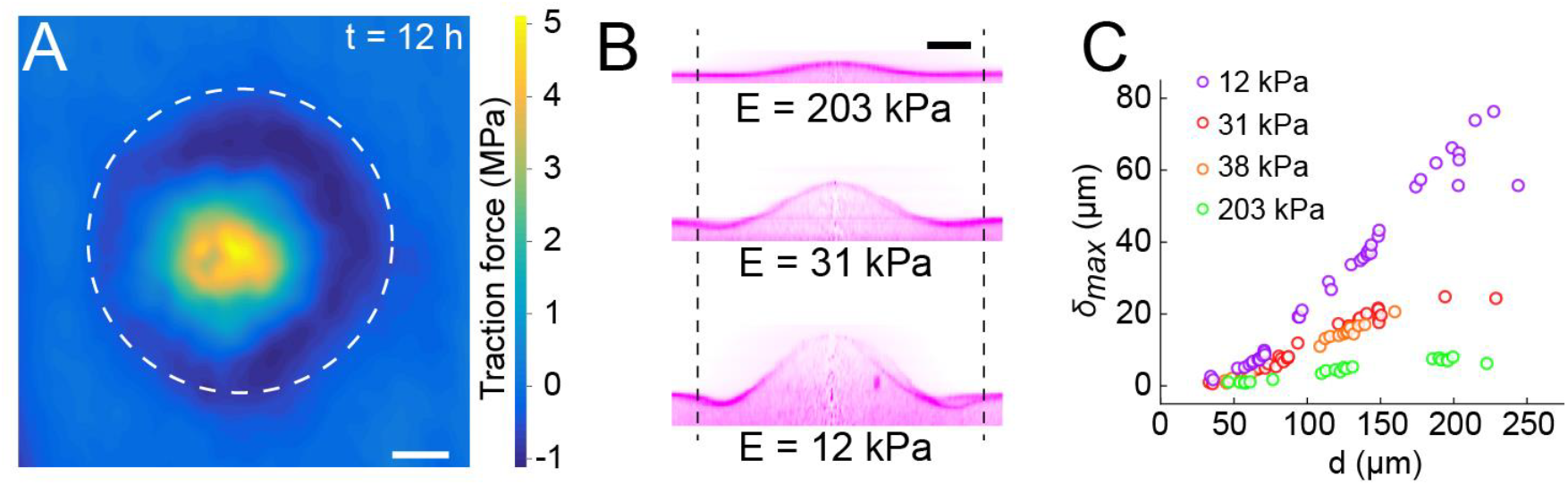
Biofilms generate large traction forces. (A) Traction force microscopy measurements at the hydrogel-biofilm interface. The dashed line shows the edge of the biofilm. Traction force is largest at the biofilm center, reaching 5 MPa. (B) Deformation profiles generated by *V. cholerae* biofilms of equal diameters on three hydrogels with different stiffness. (C) Biofilm diameter-dependence of maximum deformation for four different hydrogel composition representing a typical range of tissue stiffnesses. The softest hydrogel can deform up to 80 µm for a biofilm diameter of 220 µm. Scale bar: 20 µm.

Given the large forces generated by biofilms on hydrogel substrates, we wondered to which extent they could deform biomaterials of different stiffnesses as defined by their Young’s modulus. To test this, we reproduced the mechanical properties of various tissue types by tuning the stiffness of the PEG hydrogel films between 10 kPa and 200 kPa (24, 25). The stiffest hydrogels only slightly deformed (Fig. 4*B*, *δ*_*max*_ = 5 µm for *E* = 203 kPa). In contrast, biofilms growing on the softest hydrogels displayed large deformations (*δ*_*max*_ = 27 µm for *E* = 12 kPa). The rate of increase of deformations was inversely correlated with stiffness, resulting in differences in *δ*_*max*_ between colonies of identical diameter growing on substrates with distinct stiffnesses (Fig. 4*C*). For each stiffness, the deformation amplitude *δ*_*max*_ and the width *λ* increased linearly with biofilm diameter (Fig. 4*C* and Fig. S4). Rescaling *δ*_*max*_ with the biofilm diameter highlights a power-law relationship between deformation and substrate stiffness qualitatively consistent with the theory of buckling of plates coupled to an elastic foundation (Fig. S5)(26).

### Biofilms deform and disrupt epithelial cell monolayers

Given the ability of biofilms to generate large forces and to deform materials across a wide stiffness range, we wondered whether they could disrupt soft epithelium-like tissues. To test how biofilms can mechanically perturb host tissue during colonization, we engineered epithelial cell monolayers at the surface of soft extracellular matrix. Such cell-culture system replicates the mechanical properties of host epithelia including tissue stiffness and adhesion to underlying ECM. As a result, it constitutes a more realistic host-like environment compared to cell monolayers grown on plastic or glass. We thus engineered epithelial monolayers of enterocyte-like Caco-2 cells on a soft extracellular matrix composed of Matrigel and collagen (Fig. 5*A*). This produced soft and tight ECM-adherent epithelia. We seeded the surface of these epithelia with *V. cholerae* WT*. We note that the WT* strain has reduced virulence compared to WT *V. cholerae* due to its constitutively high levels of cyclic-di-GMP which decreases the expression of virulence factors to promote the biofilm state (27). *V. cholerae* biofilms formed at the epithelial surface within 20 h (Fig. 5*B*). Overall, biofilms perturbed the shape of the epithelium. Under biofilms, the cell monolayer detached from its ECM substrate and was often bent as did synthetic hydrogel films (Fig. 5*B-ii*). More surprisingly, we also observed that Caco-2 cell monolayers lost cohesion and single cells were engulfed by the biofilm. This allowed the biofilm to breach the epithelium and reach the ECM. There, biofilms deformed the ECM substrate, turning the initially flat surface into a dome-like shape as our synthetic hydrogels did (Fig. 5*B-iv*). These disruptions did not depend on host cell type as *V. cholerae* could also damage and bend monolayers of MDCK cells which has strong cell-cell junctions (Fig. 5*C*) (28). Our observations suggest that biofilms apply mechanical forces on host tissue thereby perturbing the morphology and integrity of epithelia, as well as its underlying ECM.

**Fig. 5:**
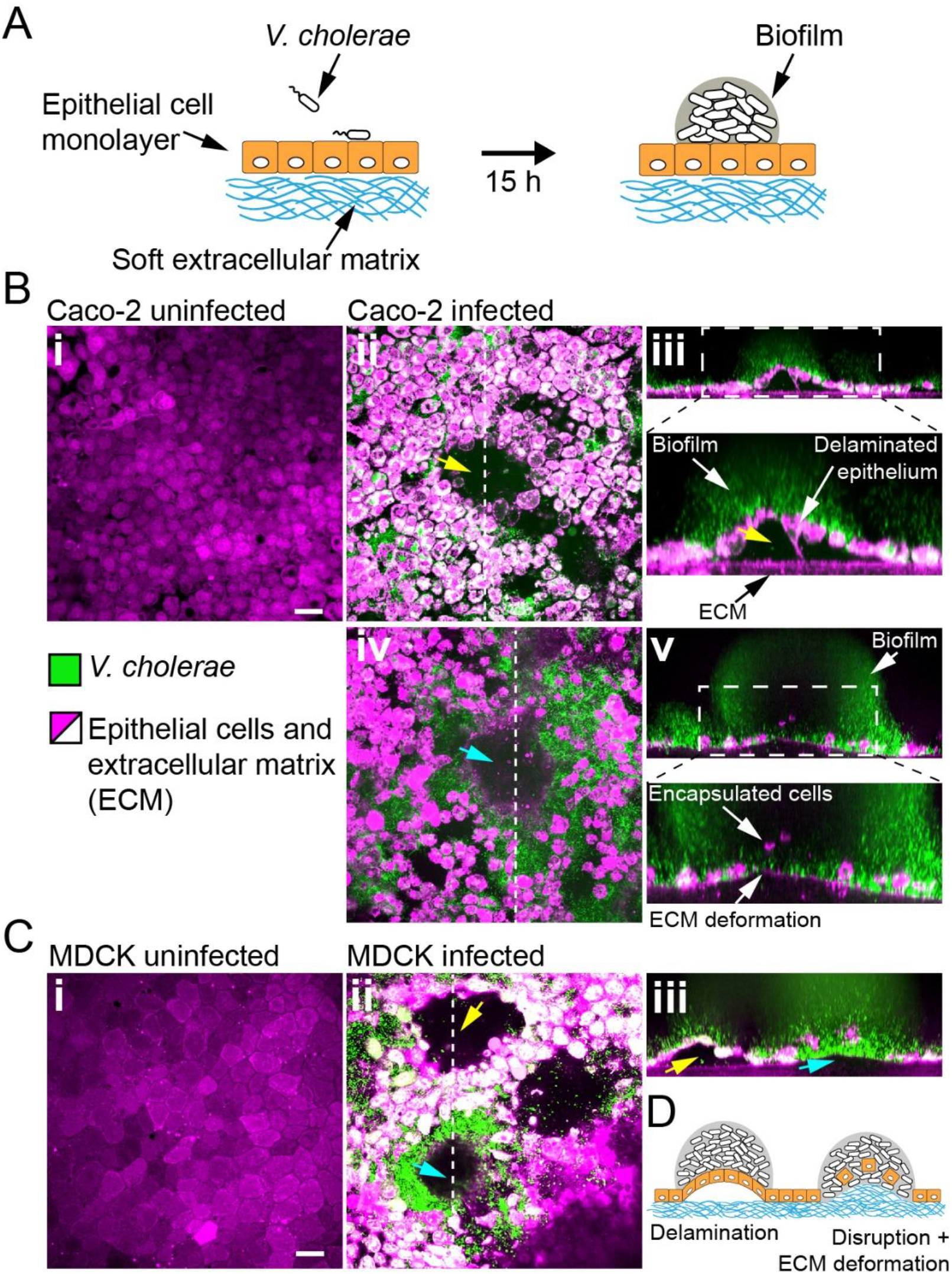
Biofilms deform and disrupt epithelial cell monolayer. (A) Caco-2 and MDCK cells grow at the surface of a soft ECM into a tight monolayer on which we seed a liquid inoculum of *V. cholerae*. (B) Confocal images of uninfected (i) and infected (ii-v) monolayers of Caco-2 cells. Yellow arrow indicates gaps in the epithelial monolayer (ii and iii), blue arrow shows deformed ECM (iv). (C) Confocal images of uninfected (i) and infected (ii-iii) monolayers of MDCK cells, also showing delamination and rupture as illustrated in (D). Scale bars: 20 µm.

## Discussion

We demonstrated that biofilms can deform the surface of soft materials they grow on. We observed that both *V. cholerae* and *P. aeruginosa* generate these deformations, suggesting that it is a feature of biofilm growth and is not species-dependent. We identified key physical and biological components that enable these deformations. In particular, our measurements of hydrogel deformations provide evidence consistent with a mechanism where the biofilm buckles as it develops. This mechanism is reminiscent of Euler buckling where the internal compressive stress in a beam triggers an instability that induces transverse deformations. In our case, we found that the onset of the buckling instability depends on growth under mechanical constraint which generates a buildup of compressive stress. In-plane hydrogel strain measurements indicate a friction between the surface and the expanding biofilm, which promotes buildup of internal stress. Also, the fact that biofilms of the *V. cholerae rbmA* and *P. aeruginosa* EPS genes deletion mutants have reduced or abolished ability to buckle or to deform the surface indicates that cell-cell cohesion in the biofilm may also participate in mechanical constraint. Without cell-cell cohesion and matrix elastic property, the viscous biofilm would flow, dissipating mechanical stress and eluding the elastic instability.

These two contributions, biofilm-surface friction and matrix elasticity, induce a buildup of compressive stress within the biofilm, ultimately causing buckling. The facts that the onset of deformation occurs at a finite critical biofilm diameter and that the width of the deformation *λ* scales linearly with this diameter are consistent with an Euler-type buckling instability (29). Also, the slight negative deformations (recess) observed near the edge of larger biofilms is reminiscent of higher order buckling modes. Finally, the absence of hydrogel deformations in biofilms from the *V. cholerae bap1* mutant shows that adhesion helps transmit the transversal forces (normal to the surface) generated during buckling to the hydrogel. In addition, the fact that for stiffer substrates *V.cholerae* and *P.aeruginosa* respectively deform and delaminate from the substrate demonstrate the important balance between adhesion and substrate elasticity in this phenomenon (30). Thus, biofilms mechanically shape their environment via a buckling-adhesion mechanism, reminiscent of the buckling and wrinkling of plates and films on elastic foundations (26).

Internal stress generated by bacterial expansion under physical constrains influences the morphologies of colony biofilms, forming wrinkles, folds and blisters. These colonies shapes are also caused by a buckling/wrinkling-like instability which depends on the mechanical properties of the matrix. These mechanically-generated shapes have been observed in *V. cholerae*, *P. aeruginosa, Bacillus subtilis* and *E. coli* and have been instrumental as an obvious phenotype to identify components and regulators of the biofilm matrix and to characterize the mechanics driving multicellular growth (31–35). However, the impact of these macroscale morphological changes and internal mechanics on the physiology of resident microbes have yet to be identified. Immersed, micrometer scale biofilms that are commonly found in natural microbial niches also undergo architectural transitions due to the emergence of internal mechanical stress. For example, cell-cell cohesion coupled with growth participates in the alignment of single cells within the multicellular structure (36). In addition, a buckling instability causes *V. cholerae* cell verticalization in the initial step of biofilm formation, in a mechanism that depends on friction of single cells with their glass substrate, generating compressive mechanical stress (14). Single cells in *E. coli* microcolonies reorient through a similar mechanism (15). The physiological functions of these cellular rearrangements have however not yet been identified. The buckling-adhesion model we here propose is consistent with the mechanics of immersed and colony biofilms. Our observations suggest that internal mechanical stress can have a function in the interaction between the biofilm and its surrounding environment, influence the morphology and mechanics of its material substrate. This may result in fouling of abiotic surfaces, in damaging competing biofilms or even host tissues.

Despite being widespread in the environments of microbes, the influence of substrate rigidity is generally overlooked in studies of surface attachment and biofilm formation (37–39). Using a materials approach aimed at reproducing a host-like environment, we found that substrate mechanical properties have a strong impact on biofilm development. Biofilm-induced deformations are particularly relevant when considering their growth at the surface of soft biological tissues. We demonstrated that biofilms generate large forces, and that these forces can be transmitted to underlying epithelia. In response, we observed that epithelial monolayers delaminate from their ECM and subsequently bend. The biofilm-generated forces also disrupt epithelial monolayers. Consistent with this, traction force microscopy measurements show that biofilms can generate MPa surface stress, which is larger than the strength of epithelial cell-cell junctions that typically rupture under the kPa range (40). In summary, our visualizations in tissue-engineered epithelia and on hydrogel films suggest that biofilms could mechanically damage host tissues when growing *in vivo*. Consistent with this hypothesis, many biofilms are known to cause tissue lesions. For example, the urine of vaginosis patients contains desquamated epithelial cells covered with biofilms (41, 42). Commensal biofilms form scabs at the epithelial surface of honeybee’s gut, triggering immune responses (43). Epithelial integrity is also compromised in intestinal diseases such as inflammatory bowel disease in a process that highly depends on the composition of the microbiota (44). Finally, hyper-biofilm forming clinical variants of *P. aeruginosa* cause significant damage to the surrounding host tissue despite its reduced virulence (45). Mechanical interactions between bacterial collectives and their host may thus represent an overlooked contributor of infections, colonization and dysbiosis. Further investigations will address whether non-pathogenic biofilm-forming species can induce epithelial damage and in fact contribute to chronic inflammation.

Most studies of biofilm formation have so far focused on their internal organization and mechanics and on the genetic regulation of matrix production. How biofilms physically interact with their natural environments has been however vastly unexplored, but is necessary knowledge to generate a holistic understanding of host-microbe interactions. This will require the development of innovative techniques that can reproduce physical components of the natural environments of biofilm-forming species in the lab such as the ones presented here.

## Acknowledgements

We would like to thank the Fitnat Yildiz, Matt Parsek, Melanie Blokesch and Bonnie Bassler for strains and plasmids and John Kolinski and Pedro Reis for discussions.

## Methods

### Cell culture

Caco-2 cells and MDCK cells were maintained in T25 tissue culture flasks (Falcon) with DMEM medium (Gibco) supplemented with 10% fetal bovine serum at 37°C in a CO_2_ incubator.

### Cell culture on collagen/Matrigel gels

To resemble the extracellular matrix natural niche, we cultured epithelial cells at the surface of collagen and Matrigel based hydrogels. Hydrogel solutions were prepared on ice to avoid premature gelation by mixing 750 µl of neutralized collagen with 250 µl of growth-factor reduced Matrigel matrix (Corning, 356231). The neutralized collagen was obtained by mixing 800 µl of native type I collagen isolated from the bovine dermis (5mg/ml, Cosmo Bio Co., Ltd.) with 10 µl of NaHCO_3_ (1 M), 100 µl of DMEM-FBS and 100 µl of DMEM 10X. We then spread 100 µl of the hydrogel solution in glass bottom dishes (**P35G-1.5-20-C, MatTek**), which were kept on ice. Excess solution was removed from the sides of the well to avoid the formation of a meniscus. To promote collagen adhesion, the wells were previously functionalized with a 2% polyethyleneimine solution (Sigma-Aldrich) for 10 min and a 0.4% glutaraldehyde solution (Electron Microscopy Science) for 30 min. We finally placed the coated dishes at 37°C in a CO_2_ incubator for 20 minutes to allow gelation.

MDCK and Caco-2 cells were detached from the flask using trypsin (Sigma-Aldrich). We seeded the cells at a concentration of 1000 cells/mm^2^ on top of the gels. We let the cells adhere for 1 day and then we filled the dishes with 2 ml of culture medium. The medium was changed every 2 days.

### Bacterial strains and culture conditions

A list of the strains and plasmids is provided in Table S1. All strains were grown in LB medium at 37°C. Deletion of the *V. cholerae* genes *rbmA* and *bap1* were generated by mating a parental A1552 *V. cholerae* strain, rugose variant, with *E. coli* S17 strains harboring the deletion constructs according to previously published protocols (46). *P.aeruginosa* strains (PAO1 parental strain) are all constitutively expressing GFP (attTn7::miniTn7T2.1-Gm-GW::PA1/04/03::GFP).

### Infection of tissue-engineered epithelia by *Vibrio cholerae*

*V. cholerae* was grown in LB medium at 37°C to mid-exponential phase (OD 0.3-0.6). Bacteria were washed 3 times by centrifugation and resuspension in Dulbecco’s phosphate-buffered saline (D-PBS). The cultures were then diluted to an optical density of 10^−7^ and filtered (5.00 µm-pore size filters, Millex) to ensure the removal of large bacterial clumps, thereby isolating planktonic cells. This ensured that biofilms growing on epithelia formed from single cells. We loaded 200 µL of diluted culture on top of Caco-2 or MDCK cells that were cultured for 1 to 7 days post-confluence on collagen/Matrigel gels. Bacteria were allowed to adhere to the surface for 20 minutes, after which cells were rinsed two times with D-PBS.

For the implementation of the flow on top of Caco-2 cells, we prepared a circular slab of PDMS with the same dimensions as the dish. We punched 1mm inlet and outlet ports in this PDMS slab. We then glued it to the rim of the dish, where no cells are present. We then connected the inlet port to a disposable syringe (BD Plastipak) filled with culture medium using a 1.09 mm outer diameter polyethylene tube (Instech) and a 27G blunt needle (Instech). The syringes were mounted onto a syringe pump (KD Scientific) positioned inside a CO_2_ incubator at 37°C. The volume flow rate was set to 50 µL⋅min^−1^.

For stationary biofilm growth on MDCK cells, the glass bottom dishes were filled with 2 mL of culture medium and were incubated at 37°C in a CO_2_ incubator.

### Fabrication of PEG hydrogels and mechanical characterization

To generate PEG hydrogels films we prepared solutions of M9 minimal medium containing poly(ethylene glycol) diacrylate (PEGDA) as the precursor and lithium phenyl-2,4,6-trimethylbenzoylphosphinate (LAP, Tokio Chemical Industries) as the photoinitiator. Molecular weight and concentration of PEGDA were tuned to obtain hydrogels with different stiffnesses (Table S2), while the concentration of LAP is kept constant at 2 mM. To incorporate fluorescent microparticles into the PEG hydrogels, we modified the original solution by substituting 2 µL of M9 medium with 2 µL of red fluorescent particles solution (ThermoFischer, FluoSpheres, Carboxylate-modified Microspheres, 0.1 µm diameter, 2% solids, F8887).

To prepare the samples for mechanical characterization, we filled PDMS wells (5 mm diameter, 4 mm height) with the hydrogel solution. We covered the wells with a coverslip and we let them polymerize in a UV transilluminator (Bio-Rad Universal Hood II) for 5 minutes. The resulting hydrogel cylinders were immersed in M9 overnight and tested with a rheometer (TA instruments) in compression mode, at a deformation rate of 10 µm/s. Beforehand, the diameter of the cylinders was measured with a digital caliper, while the height of the cylinder was defined as the gap distance at which the force starts differing from zero. The elastic modulus corresponds to the slope of the linear fit of the stress-strain curves in the range of 15% strain. The final modulus is the average modulus of 3 replicates.

### Fabrication of thin PEG hydrogel layers and implementation with PDMS microfluidic chip

We fabricated microfluidic chips following standard soft lithography techniques. More specifically, we designed 2 cm-long, 2 mm-wide channels in Autodesk AutoCAD and printed them on a soft plastic photomask. We then coated silicon wafers with photoresist (SU8 2150, Microchem), with a thickness of 350 µm. The wafer was exposed to UV light through the mask and developed in PGMEA (Sigma-Aldrich) in order to produce a mold. PDMS (Sylgard 184, Dow Corning) was subsequently casted on the mold and cured at 70 °C overnight. After cutting out the chips, we punched 1 mm inlet and outlet ports. We finally punched a 3 mm hole right downstream of the inlet port. This hole, after being covered with a PDMS piece, acts as a bubble trap.

To obtain thin and flat hydrogel layers, a drop of about 80 µL of the hydrogel solution was sandwiched between two coverslips and incubated in the UV transilluminator for 5 minutes to allow gelation. The bottom coverslip (25×60 mm Menzel Gläser) was cleaned with isopropanol and MilliQ water, while the upper one (22×40 mm Marienfeld) was functionalized with 3-(Trimethoxysilyl)propyl methacrylate (Sigma-Aldrich) following the standard procedure. In short, cleaned coverslips were immersed in a 200 mL solution of ethanol containing 1 mL of the reagent and 6ml of dilute acetic acid (1:10 glacial acetic acid:water) for 5 minutes. They were subsequently rinsed in ethanol and dried. This functionalization enables the covalent linkage of the hydrogel to the coverslip.

Right after polymerization, the coverslips were separated using a scalpel and thus exposing the hydrogel film surface. We then positioned the PDMS microfluidic chip on top of the hydrogel film. This results in a reversible, but sufficiently strong bond between the hydrogel and the PDMS, allowing us to use the chips under flow without leakage for several days. The assembled chips were filled with M9 to maintain the hydrogel hydrated.

### Biofilm growth in microfluidic chambers

All *V. cholerae* and *P. aeruginosa* strains were grown in LB medium at 37°C until mid-exponential phase (OD 0.3-0.6). The cultures were diluted to an optical density of 10^−3^ and subsequently filtered (5.00 µm-pore size filters, Millex) to ensure the removal of large bacterial clumps. We then loaded 6.5 µL of the diluted bacterial culture in the channels, from the outlet port. We let them adhere for 20 minutes before starting the flow. We connected the inlet port to a disposable LB-filled syringe (BD Plastipak) mounted onto a syringe pump (KD Scientific), using a 1.09 mm outer diameter polyethylene tube (Instech) and a 27G needle (Instech). For all conditions, the volume flow rate was 10 µL⋅min^−1^, which corresponds to a mean flow speed of about 0.25 mm⋅s^−1^ inside the channels. The biofilms were grown at 25°C.

### Staining procedures

Caco-2 cells and MDCK cells were incubated for 20 minutes in a 10 µM solution of CellTracker Orange CMRA (Invitrogen, C34551) and washed with DPBS before seeding the bacteria.

Since *V. cholerae* strains were not constitutively fluorescent, biofilms were incubated for 20 minutes with a 10 µM solution of SYTO9 (Invitrogen, S34854) and washed with M9 minimal medium before visualization. This results in double staining of epithelial cells in the case of infection experiments.

### Visualization

For all visualizations, we used an Nikon Eclipse Ti2-E inverted microscope coupled with a Yokogawa CSU W2 confocal spinning disk unit and equipped with a Prime 95B sCMOS camera (Photometrics). For low magnification images, we used a 20x water immersion objective with N.A. of 0.95, while for all the others we used a 60x water immersion objective with a N.A. of 1.20. We used Imaris (Bitplane) for three-dimensional rendering of z-stack pictures and Fiji for the display of all the other images.

To obtain the deformation profiles, z-stacks of the hydrogel containing fluorescent microparticles were performed every 0.5 µm, while a brightfield image of the base of the biofilm was taken to allow measurement of the diameter of the biofilm. For the visualization of the full biofilm, z-stacks of the samples were taken every 2-3 µm. For timelapse experiments, biofilms were imaged as soon as the flow was started, while for all the other experiments biofilms were imaged between 10 and 24 h post-seeding.

### Image analysis and computation of deformation profiles

Starting from confocal imaging pictures of the microparticle-containing hydrogel, we aimed at identifying the gel surface and extracting quantitative information about its deformation induced by the biofilms. In most cases, we used an automated data analysis pipeline as described below. To get an average profile of the deformation caused by the biofilms, we performed a radial reslice in Fiji over 180 degrees around the center of the deformation (one degree per slice). We then performed an average intensity projection of the obtained stack. Tocalculate the diameter of the biofilm, we averaged 4 measurements of the biofilm diameter taken at different angles. The resliced images were then imported in Matlab R2017a (Mathworks) as two-dimensional (x-y) matrices of intensities. In these images, the surface was consistently brighter than the rest of the gel. Therefore, we identified the surface profile as the pixels having the maximal intensity in each column of the matrix. Note that the bottom of the gel sometimes also comprised bright pixels that introduced noise in the profile. To reduce this problem, we thus excluded 20 rows at the bottom of each image (~3.7 µm). We then calculated the baseline position of our gel – namely, the height of the non-deformed portion of the gel. In our pictures, this corresponds to the height at the left and right extremities of the profile. Therefore, we defined the baseline as the average of the first 50 and last 50 pixels of the profile (~9 µm on each side of the profile). We then offset the whole picture so that the baseline position corresponded to y = 0. We undersampled the extracted surface profiles to further reduce noise, by keeping only the maximal y value over windows of 40 pixels. Finally, we fitted a smoothing spline to the undersampled profile using the built-in *fit* function in Matlab, with a smoothing parameter value of 0.9999.

To quantify the deformation that biofilms induced on the hydrogel, we measured the amplitude (*δ*_*max*_) of the deformed peak and its full width at half maximum (λ). First, we evaluated the fitted profile described above at a range of points spanning the whole width of the picture and spaced by 0.0005 µm. We identified the maximal value of the profile at these points, which corresponds to the amplitude of the peak *δ*_*max*_ (with respect to the baseline, which is defined as y = 0). We then split the profile in two: one part on the left of the maximum, and one part on its right. On each side, we found the point on the profile whose y value was the closest to 0.5 ⋅ *δ*_*max*_ using the Matlab function *knnsearch*. We then calculated the distance between their respective x values, which corresponds to the λ of the deformed peak. Our data analysis program also included a quality control feature, which prompted the user to accept or reject the computed parameters. When imaging quality was insufficient to ensure proper quantification with our automated pipeline, we measured the deformation manually in Fiji.

### Digital volume correlation and traction force microscopy

We performed particle tracking to measure local deformations and ultimately compute stress and traction forces within hydrogels as biofilms grew. To do this, we performed timelapse visualizations of the hydrogel during the formation of a biofilm at high spatial resolution with a 60X, NA 0.95 water immersion objective. We thus generated 200 µm x 200 µm x 25 µm (50 stacks of 1200×1200 pixels) volumes at 14 different time points. These images were subsequently registered to eliminate drift using the Correct 3D Drift function in Fiji. To compute local material deformations which we anticipated to generate large strains, we used an iterative Digital Volume Correlation (DVC) scheme (16). These were performed with 128×128×64 voxel size in cumulative mode, meaning deformations are calculated by iterations between each time point over the whole 4D timelapse, rather than directly from the reference initial image. The DVC code computes material deformation fields in 3D which we subsequently use as input for the associated large deformation traction force microscopy (TFM) algorithm (16). The TFM calculates stress and strain fields given the material’s Young modulus (*E* = 38 kPa in our case) to ultimately generate a traction force map at the hydrogel surface.

---

**Table S1.** Plasmids and strains used in this study

| Strain or plasmid | Relevant genotype | Source |
| --- | --- | --- |
| pFY_113 | plasmid for generation of in-frame *rbmA* deletion mutants | (47) |
| pFY_330 | plasmid for generation of in-frame *bap1* deletion mutants | (47) |
| *V. cholerae* O1 El Tor A1552 (*Vc* WT*) | rugose variant | (48) |
| *V. cholerae* Δ*rbmA* | in frame deletion of *Rbma* in rugose backgrounds | This study |
| *V. cholerae* Δ*bap1* | in frame deletion of *Bap1* in rugose backgrounds | This study |
| PAO1 WT | wild-type, *Gm*<sup>r</sup> | (49) |
| PAO1Δ*wspF* (PAO1 WT*) | in frame deletions of *WspF*, *Gm*<sup>r</sup> | (50) |
| PAO1Δ*wspF*Δ*pel* | in frame deletions of *WspF*, *PelA* genes, *Gm*<sup>r</sup> | (50) |
| PAO1Δ*wspF*Δ*psl* | in frame deletions of *WspF*, *PsIBCD* genes, *Gm*<sup>r</sup> | (50) |
| PAO1Δ*wspF*Δ*cdrA* | in frame deletions of *WspF*, *PsIBCD*, *cdrA* genes, *Gm*<sup>r</sup> | (51) |

**Table S2.** Molecular weight and concentrations of the precursors used for the generation of the hydrogels and resulting elastic modulus

| Precursor | Concentration wt/vol | Modulus kPa |
| --- | --- | --- |
| PEGDA MW 10000 (Biochempeg) | 10% | 12.1 ± 0.8 |
| PEGDA MW 6000 (Biochempeg) | 10% | 38.3 ± 1.0 |
| PEGDA MW 3400 (Biochempeg) | 10% | 30.9 ± 2.0 |
| PEGDA MW 700 (Sigma-Aldrich) | 15% | 203.3 ± 13.7 |

