## Extended data for "Biofilms deform soft surfaces and disrupt epithelia"

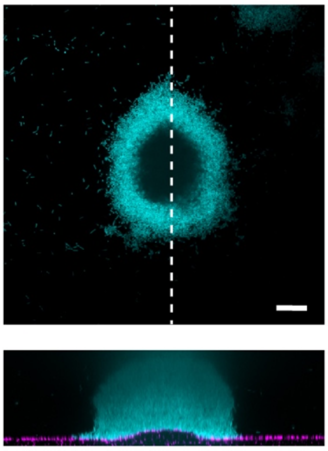

**Fig. S1.** Deformation of hydrogel substrate (E = 38 kPa) by biofilms of PAO1 WT. Scale bar: 20 µm

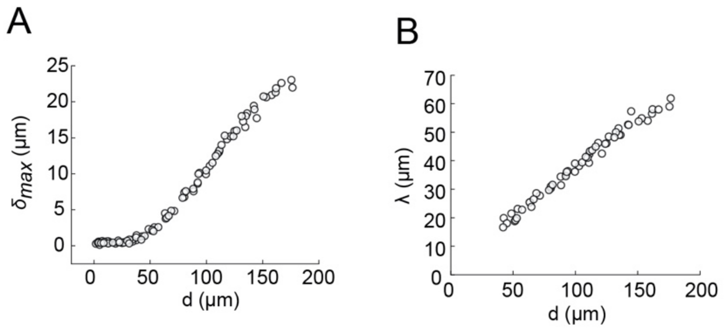


**Fig. S2. (**A) Biofilm diameter-dependence of $\delta_{max}$. (B) Biofilm diameter-dependence of *λ.* Both $\delta_{max}$ and *λ* linearly scale with the diameter *d* of the biofilm.

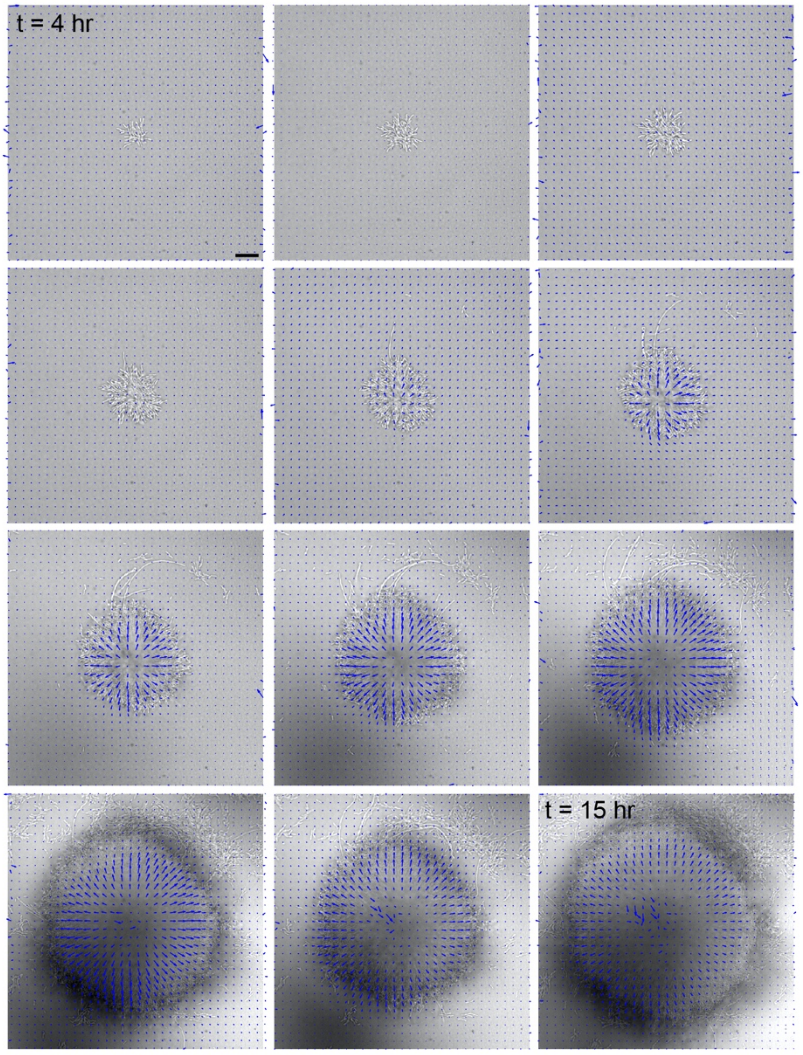


**Fig. S3.** Hydrogel deformation field computed at different growth stages, superimposed with a brightfield image of the biofilm. Scale bar: 20 µm. The force field at each timestep is normalized by its maximum displacement, thereby showing relative deformations.

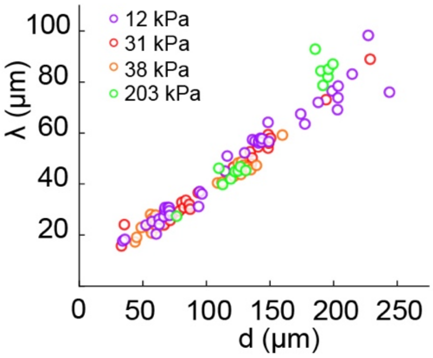


**Fig. S4.** Biofilm diameter-dependence of *λ* for substrates with different moduli. *λ* scales linearly and it is not substrate-stiffness dependent.

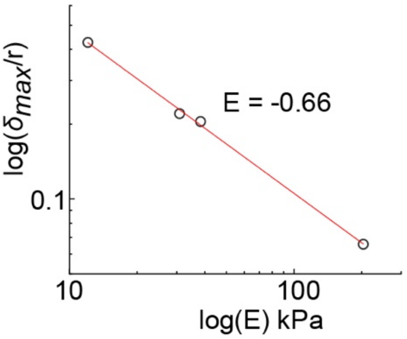


**Fig. S5.** Power-law relationship between deformation $\delta_{max}$ and substrate moduli (E).

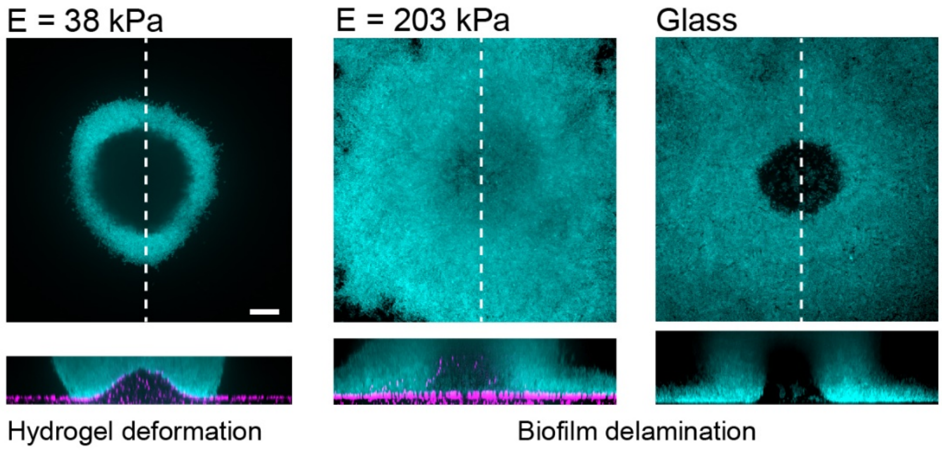


**Fig. S6.** *P. aeruginosa* biofilms on substrates with different stiffness. Increasing hydrogel stiffness to 200 kPa induces delamination of biofilms, as observed on glass. Scale bars: 20 µm.

**Movie S1.** Timelapse visualization of *V. cholerae* WT* biofilm growth (brightfield) and corresponding hydrogel deformation (*E* = 38 kPa) in the *xy* and *xz* planes. Scale bar 20 μm.
